# Evaluating the roles of testosterone and sex-linked genes in territorial aggression of sex-reversed XY females in the African pygmy mouse

**DOI:** 10.64898/2025.12.05.692630

**Authors:** Louise D. Heitzmann, Julie Perez, Romann Charbonnier, Marc Fichter, Xavier Hautecoeur, Theo Deremarque, Agnes O. Martin, Frederic Veyrunes

**Affiliations:** Institut des Sciences de l’Evolution de Montpellier, University of Montpellier, CNRS, IRD, UMR 5554, Montpellier, France; Department of Evolution, Ecology and Organismal biology, University of California, Riverside, USA; Institut de Génétique Humaine, University of Montpellier, CNRS, UMR 9002, Montpellier, France; Institut de Génétique Fonctionnelle, University of Montpellier, CNRS, INSERM, UMR 5203, Montpellier, France

**Keywords:** Territoriality, sex chromosomes, androgen, neuropeptides, dopamine

## Abstract

Sex differences in social aggression are widespread across the animal kingdom, with males typically displaying greater territoriality. While this dimorphism has traditionally been attributed to sex hormones, sex chromosomes can also contribute to it independently of hormonal influence. In the African pygmy mouse *Mus minutoides*, naturally occurring sex-reversed XY females (named X*Y due to a mutation on the X chromosome) are highly territorial in comparison to the other female genotypes present in the population (XX and XX*). However, the molecular basis of this phenotype remains unknown. Here, we evaluate molecular factors – known to correlate with aggressiveness – following a standardized behavioural assessment of aggression. We focus on i) the androgen pathway by quantifying testosterone serum levels and expression of its receptor in the brain; ii) the brain dopaminergic system through expression of dopamine-regulating genes, including the sex-determining *Sry* gene; and iii) neuroendocrine circuits, via vasopressin and oxytocin expression. These systems, although distinct, converge on hypothalamic-limbic circuits that regulate social behaviour and threat responses, providing an integrated framework for the modulation of aggression. Contrary to earlier reports, we found reduced level of aggression in X*Y females, likely to reflect a breeding facility effect. We also observe no correlations between aggressiveness and androgen levels or gene expression of the tested factors. However, our results support a stimulation of the dopaminergic system and of the oxytocin pathway following the agonistic assay suggesting their potential involvement in aggression-related responses. This further supports the idea that aggression is multifactorial. It is shaped by the interaction of several neuroendocrine and neurotransmitter pathways rather than a single determinant.

## Introduction

Aggression is a complex social behaviour widespread across the animal kingdom. In many species, social aggression is highly sexually dimorphic, prevalent in males competing for territories and mates, whereas female aggression is often associated with maternal defense (Darwin, 1871; Pandolfi et al. 2021). This sexual dimorphism is mostly known to arise from sex differences in steroid hormones levels, especially testosterone which can promote aggression either directly or through its metabolites (e.g. aromatization of testosterone to estradiol) (Rose et al. 1971; Book et al. 2001; Han & De Vries, 2003; Giammanco et al. 2005; Phoenix, 2009; Phoenix et al. 1959; Wu et al. 2009; Zha et al. 2025).

Sex chromosomes and imbalance in sex-linked genes can also contribute to sex differences in social aggression, factors that have received relatively little attention (Gatewood et al. 2006; Lovell-Badge, 2005; Saunders et al. 2016; Sluyter et al. 1996). For instance, levels of dopamine – a well-characterized neurotransmitter associated with both social and maternal aggression – can be directly influenced by the expression of sex-linked genes (Czech et al. 2012; Godar et al. 2016; Korzan et al. 2006; Kudryavtseva et al. 2017; Mahadevia et al. 2021; van Erp & Miczek, 2000; Yamaguchi & Lin, 2018). The sex-determining Y-linked gene *Sry* is expressed in specific brain regions and upregulates the rate-limiting enzyme required for dopamine production: tyrosine hydroxylase (*Th*). Elevated level of *Th,* and associated dopamine levels, is directly correlated to increased agonistic behaviours (Mahadevia et al. 2021). Conversely, the gene encoding the enzyme responsible for dopamine degradation, monoamine oxydase A enzyme (*MaoA;* Shih et al. 1999), is located on the X chromosome. Consistent with the relationship between dopamine levels and aggression, knock-out or reduced expression of *MaoA* have been related to enhanced aggressiveness in mice (Cases et al. 1995; Scott et al. 2008; Kudryavtseva et al. 2017) and humans (Godar et al. 2016;). There is also evidence that the neuropeptide vasopressin (AVP), which can promote or inhibit aggression (depending on species), is influenced by X chromosome number (Gatewood et al. 2006; Bester-Meredith & Marler, 1999; de Moura Oliveira et al. 2021; Everts et al. 1997; Cox et al. 2015). These examples highlight that sex-linked genes (and the autosomal genes they regulate) can shape the neural circuits underpinning aggressive behaviours, but their role is often difficult to separate from that of steroid hormones as chromosomal sex (XX or XY) usually aligns with gonadal sex (ovaries or testes). Nonetheless, there are a few mammalian exceptions in which these two factors are dissociated (Saunders & Veyrunes, 2021).

The African pygmy mouse is a mammal species showing a natural dissociation between gonadal and chromosomal sexes, with the presence of sex-reversed females. About 1 million years ago, there was a mutation on the X chromosome resulting in a third sex chromosome, called X*, and induced sex reversal in X*Y individuals (Veyrunes et al. 2010a). Three female genotypes are found in natural populations: XX, XX* and X*Y, while all males are XY. In addition, the three sex chromosomes are fused to autosomes, so called neo-sex chromosomes, which are subject to sex-specific recombination and inheritance patterns as well (Veyrunes et al. 2010b; Veyrunes et al. 2006; Baudat et al. 2019; Saunders et al. unpublished data). Although X*Y individuals are fully fertile females (Rahmoun et al. 2014; Saunders et al. 2014), they have different behavioural phenotypes compared to XX and XX* females, including an enhanced territoriality assessed by aggressive behaviours in a resident-intruder assay (Saunders et al. 2016). Yet, whether sex chromosomes influence directly (by regulating the expression of sex-linked and/or autosomal genes; e.g., dopamine pathway) or indirectly (through their control on hormone levels and notably testosterone) social aggression is unknown.

Here, we present a first attempt to reproduce and conduct the agonistic assay from Saunders et al. (2016) to investigate candidates for the molecular basis of territorial aggression in X*Y females: testosterone and sex chromosomes/sex-linked genes. We evaluate agonistic behaviours from which we define a probability of offensiveness as well as an aggressiveness level (i.e., intensity of aggression) in individuals of the three female genotypes. We then correlated the aggressiveness level with markers of the androgen (hormonal output and receptor expression), dopaminergic (expression of sex-linked modulators) and neuropeptide (expression of *Avp*) pathways. Oxytocin (Oxt) was included in the neuropeptide pathway due to its established role in modulating territorial aggression as well (Tan et al. 2019; Goodson et al. 2015; Love, 2014). Our results suggest that dopamine and oxytocin pathways are stimulated following resident-intruder assays in X*Y females, but do not necessarily trigger aggressive behaviours.

## Material & Methods

### Animals

Individuals were reared in a room held at 23°C with a 15:9 light-dark cycle in our breeding facility at the University of Montpellier (CECEMA). Virgin females were kept in groups of 5-6 individuals in environmentally enriched cages (38 x 26 x 24 cm) filled with wood chip bedding, with constant food and water supply. Males were grouped by two (brothers) or isolated in small cages (32 x 45 cm) to prevent agonistic behaviour. We genotyped females by PCR of *Sry* (Veyrunes et al. 2010a) and using X/X* markers with the following primers: SC328.2.R 5’-AAGCCTTGACCCAGCCTTCT-3’ and SC328.2L1 5’-TGAAGGACAGGACAGGAACACT-3’. The primers amplified one band for XX females 553 bp, two bands for XX* females (553 and 595 bp) and one band for X*Y females (595 bp). Animal experimentations were performed with the approval of the French Ethical Committee on animal care and use (No. APAFIS#23541), and following European guidelines.

### Resident-intruder assay

Tests were conducted following the same procedures as in Saunders et al. (2016). A virgin female aged between 7-10 months (table 1) is isolated in a large plexiglass cage (40 L x 30 W x 30 H cm) with its own bedding and enrichment for 48 hours, in order to establish the resident status. We used females in the estrus stage (determined by vaginal smears with the presence of anucleate squamous-type epithelial cell; see Veryunes et al. 2024 for details) to make sure they would not be sexually receptive by the time of experimentations two days later, which is likely to influence aggression (Hyde & Sawyer, 1977). Prior the assay, male is placed in a side cage (10 x 10 x 10 cm) with a trap door leading to the experimental environment. Males are tested once or twice at most, with a minimal gap of seven days between trials. An assay started once the male entered the female environment and female-male interactions were then recorded for 10 minutes. Each experiment was recorded on camera (using phone camera) and videos were analysed manually by one rater blind to females’ genotype. We established an ethogram used by the rater to score the following behaviours: latency of the first offensive attack, frequency of attacks (i.e., number of bite attempts corrected by the overall number of female-male interactions to account for differences in female-male encounters between assays), type of attacks (defensive or offensive) as well as the percent of time females spend chasing the male during an experiment. The absence of response for each behavioural variable was set to a value of 0. At the end of each assay, females were anaesthetized and then decapitated. Blood and brain were instantaneously collected for hormonal dosage and candidate gene RNA quantifications. Given that we were interested in identifying the molecular basis underpinning differences in female territorial aggression, we did not assess male-to-male territorial aggression. Overall, we tested 19 X*Y females, 11 XX and 8 XX* females. XX and XX* females were considered as one control group with low aggression level (Saunders et al. 2016).

**Table 1.**
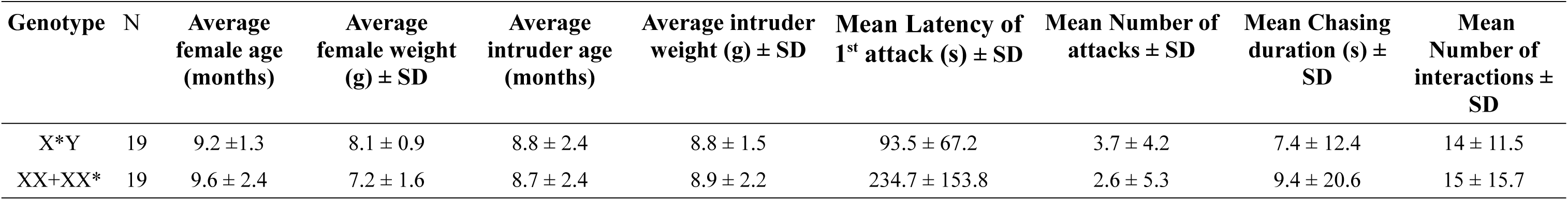
Summary statistics of females (residents) and males (intruders) during agonistic assays.

### Testosterone dosage assay

The whole blood collected after female euthanasia was left to clot for approximately 45-60 minutes. Samples were then centrifuged for 10 minutes at 4500 rpm and serum were collected into new tubes and stored at –20°C until processing. Testosterone levels were quantified by immuno-enzymatic assay using IBL International ELISA kit (#RE52151) and following manufacturer’s instructions. Females’ sera were not diluted except for two females which did not have enough material (i.e., 25 μL). These samples were treated with caution as dilution can impede detection and measures due to already low hormonal levels (Veyrunes et al. 2024). Most samples were measured only once due to limited amount of blood collected but some samples with large serum volume were duplicated and values of testosterone were then averaged across replicates. Measurement for the 38 females were spread over three ELISA assays. One female (X*Y) did not pass detection threshold and the females in the last assay (2 XX, 3 XX* and 3 X*Y) showed unusually low concentrations, indicative of unstable testosterone in serum, and likely to reflect a storage issue with prolonged thawing. We thus applied a correction factor for these values specifically: when possible (enough serum left), we repeated the ELISA test on females from the 1^st^ and 2^nd^ ELISA assays. We then calculated the difference between the first measure (used in the study) and the new ones (i.e., following thawing issue) and applied the averaged difference as a correction factor to be multiplied by the values from the 3^rd^ assay.

### RNA extraction, reverse transcription and RT-qPCR of Th, MaoA, Drd2, Sry, Avp, Oxt and Ar

Brains collected were immediately placed in RNAlater®, maintained at 4°C for 48 hours and then stored at –80 °C until further use. Whole brain RNA was extracted using Rneasy® Plus Mini Kit (Qiagen). Tissues were first grounded in liquid nitrogen using a mortar and a pestle under sterile environment. We added a lysis buffer to homogenize materials and because we used the whole tissue, we adjusted volumes to four times volumes recommendation (4 x 6 μL of β-mercapto and 4 x 600 μL of RLT buffer). This gave us four tubes per sample. RNA was purified for two tubes following manufacturer’s protocol and the remaining two were stored at –80°C. Extracted RNA was eluted in 50 μL of RNase free water and stored at –80°C until further use. Before real-time PCR quantifications (RT-qPCR) assays*, s*amples were treated with DNase (TurboDNA-free™, Invitrogen) followed by reverse transcription of 1 μg of RNA to convert into cDNA using SuperScript IV Reverse Transcriptase kit (Invitrogen), with random hexamers and following provided protocol. We then diluted cDNA samples to 1:20 prior quantifications. However, *Sry* is poorly expressed, with variable haplotypes of different lengths (Veyrunes et al. 2013), as well as in very few regions of the brain (Lahr et al. 1995; Dewing et al. 2006); its signal might be diluted and difficult to capture using whole-brain RNA and low concentrations. Therefore, for *Sry* assay specifically, we concentrated DNase-treated RNA samples (i.e.,speed vacuum centrifugation for ∼10 min at room temperature) prior the reverse transcription and diluted cDNA to 1:5 to increase RNA/cDNA yields (up to 2.5 μg of RNA) as well as qPCR performance, All RT-qPCR assays were performed on a LightCycler 480 (Roche) and using SensiFAST™ SYBR® No-ROX kit (Meridian). We used *Rps29* as a reference gene, which had previously been used (see Rahmoun et al. 2014). Expression levels of *Sry, Th*, *MaoA, Drd2, Ar, Avp* and *Oxt* were measured in triplicates and genes were distributed across three distinct assays. Primers and qPCR conditions are shown in table 2. Relative expression levels were measured on LightCycler 480 software v 1.5 (Roche). We included 12 females (4 XX, 2 XX* and 6 X*Y) that did not perform behavioural assays to control for the treatment effect on expression levels (i.e., quantification of baseline expression levels).

**Table 2.**
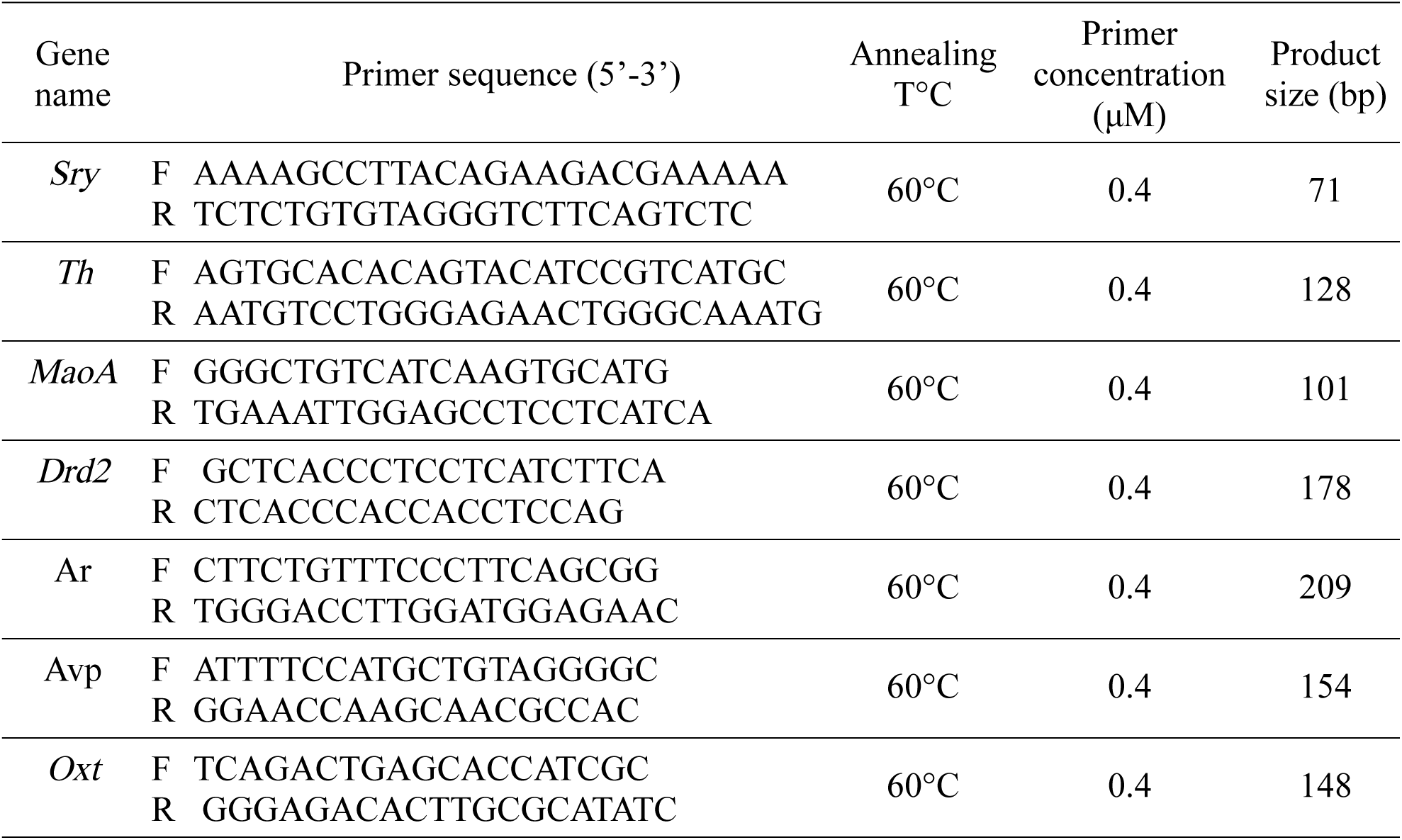
Sequences of primers and conditions for RT-qPCR.

### Statistical analyses

All statistical analyses were performed on R v4.5.2 (R core Team 2025). XX/XX* females represent the control group with low aggression level and are therefore used as the reference group for downstream analyses. We modelled the probability of offensive aggression between the two genotypic group (i.e. X*Y *vs* XX+XX* females) with a generalized linear model, fitting a binomial distribution. We then cumulated aggressive behaviours to determine an aggressiveness level: we added the frequency of attacks to the percent of time females spent chasing the male (i.e., chasing duration over the experiment duration), and to the inverse function of the latency of first attack (i.e., +1/latency). We log(x+0.1) transformed the aggressiveness level to control for heteroscedasticity and compared it between females using a linear model with a Gaussian distribution. We did not include the age nor the weight of individuals as random effects due to computational limitations (i.e., singular fit). Testosterone and gene expression levels (except for *Sry*) were log-transformed for heteroscedasticity and compared between genotypic group with linear models, fitting a Gaussian distribution. Looking at gene expression specifically, we included a treatment variable (i.e. whether females performed the behavioural assays or not: ‘Post-assay’ *vs* ‘Baseline’ categories, respectively) as a fixed effect in interaction with the genotypic group. Pairwise post-hoc comparisons were made using *emmeans* package, and p-values were adjusted for multiple testing. We then performed a Kendall rank correlation test to assess the relationship between aggressiveness level and testosterone / gene expression levels and applied a Benjamini-Hochberg correction on p-values for multiple testing. We acknowledge that the small sample size might bias our estimations (e.g low power to detect small effects) and accounted for this limitation in the discussion section accordingly.

## Results

### Agonistic behaviours

Despite a tendency for a greater aggressiveness in X*Y females (Fig. 1a-e), there is no significant difference in the probability of being offensive between X*Y females and XX/XX* females (Fig. 1d, GLM, df = 36, estimate = 0.86 ± 0.66 SE, p=0.2), nor there is a significant difference in aggressiveness level (combined agonistic behaviours, see Fig. 1, table 1) between females (Fig. 1e, LM, df = 36, estimate = 0.55 ± 0.34 SE, p=0.12).

**Figure 1.**
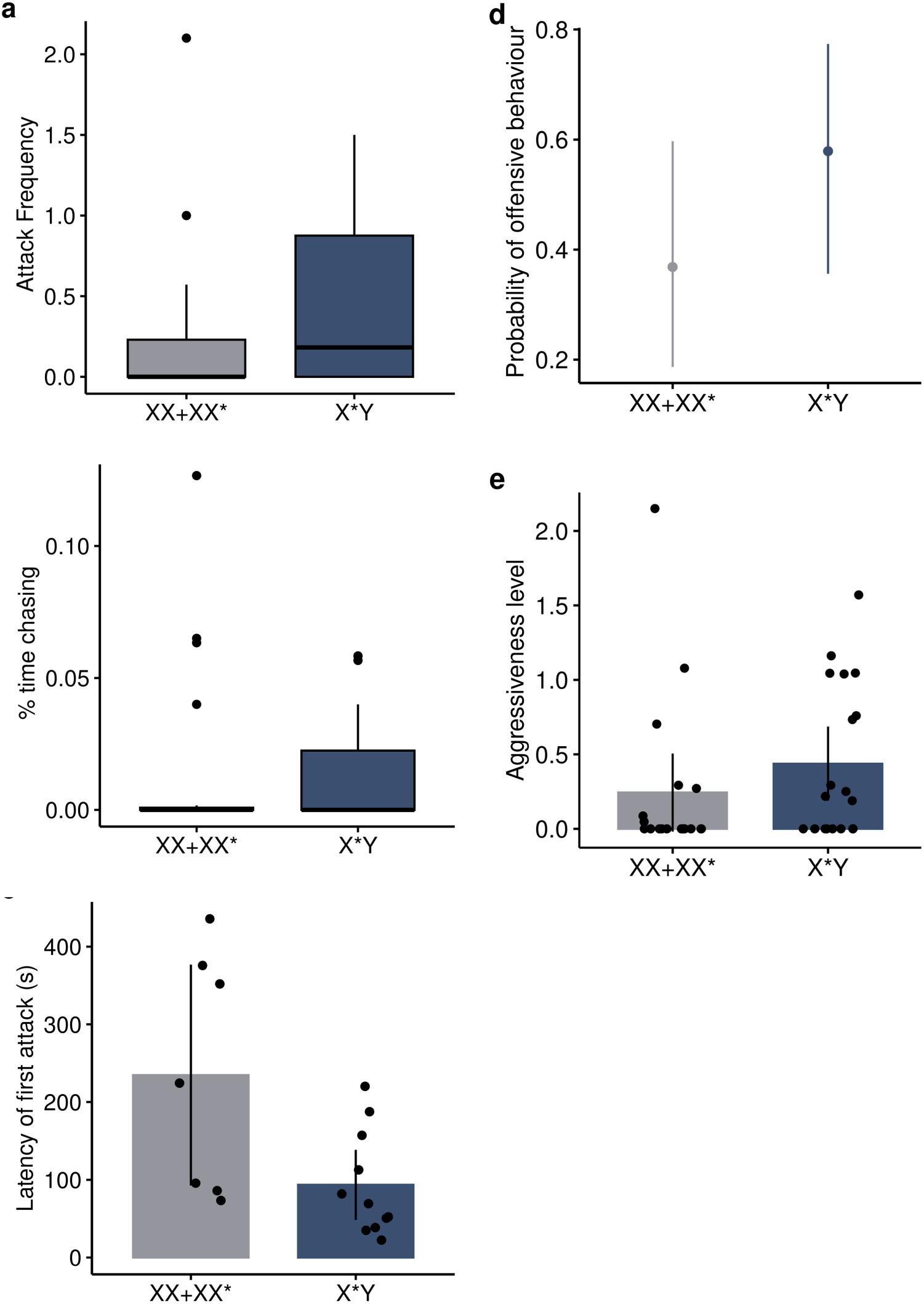
Tendency for enhanced aggressiveness in X*Y females. XX and XX* females formed the control group with low aggressiveness (Saunders et al. 2016). **a-b**) Attack Frequency (i.e. number of attacks standardized by the number of encounters) and percent of time females spent chasing males (i.e. cumulated chasing duration over the 10 min experiment), respectively. Boxplots represent median (central line), 25^th^ and 75^th^ percentile of values (box bounds). **c**) Latency of first attack (s). Values are mean ± 95 % confidence intervals. **d,** Probabilities of offensive aggression. Values are estimated average probabilities ± 95% CI. **e,** Aggressiveness level defined by the sum of agonistic behaviours (attack frequency, percent of time spent chasing and +1 for offensive behaviours). Values are mean ± 95 % confidence intervals. Dots represent raw data. n_XX+XX*_=19 and n_X*Y_ =19.

### Androgen pathway: testosterone and its receptor

There is no significant difference in average testosterone levels between X*Y females and XX/XX* females following the agonistic assay (Fig 2a, LM, df = 35, estimate = 0.16 ± 0.18 SE, p=0.38). Similarly, there are no differences in *Ar* expression levels between females, regardless of whether they were tested (Fig 2b, post-hoc Sidak, df = 46, post assay ratio = 2.03 ± 1 SE, p = 0.5,) or not (post-hoc Sidak, df = 46, baseline ratio = 1.28 ± 0.36 SE, p = 0.5). Baseline *Ar* expression levels are not significantly different from post-assay expression levels, despite a tendency for higher post-assay levels in X*Y females (post-hoc Sidak, df = 46, X*Y ratio = 0.41 ± 0.17 SE, p=0.12; XX/XX* ratio = 0.66 ± 0.26 SE, p=0.76). Finally, there is no significant correlation between androgens and aggressiveness levels in X*Y (Fig. 2c-d, Kendall test, testosterone: τ = –0.07, p=0.7; *Ar*: τ =-0.04, p=0.94) nor in XX/XX* females (Kendall test, testosterone, τ =0.31, p=0.1; *Ar*: τ =-0.007, p=0.97).

**Figure 2.**
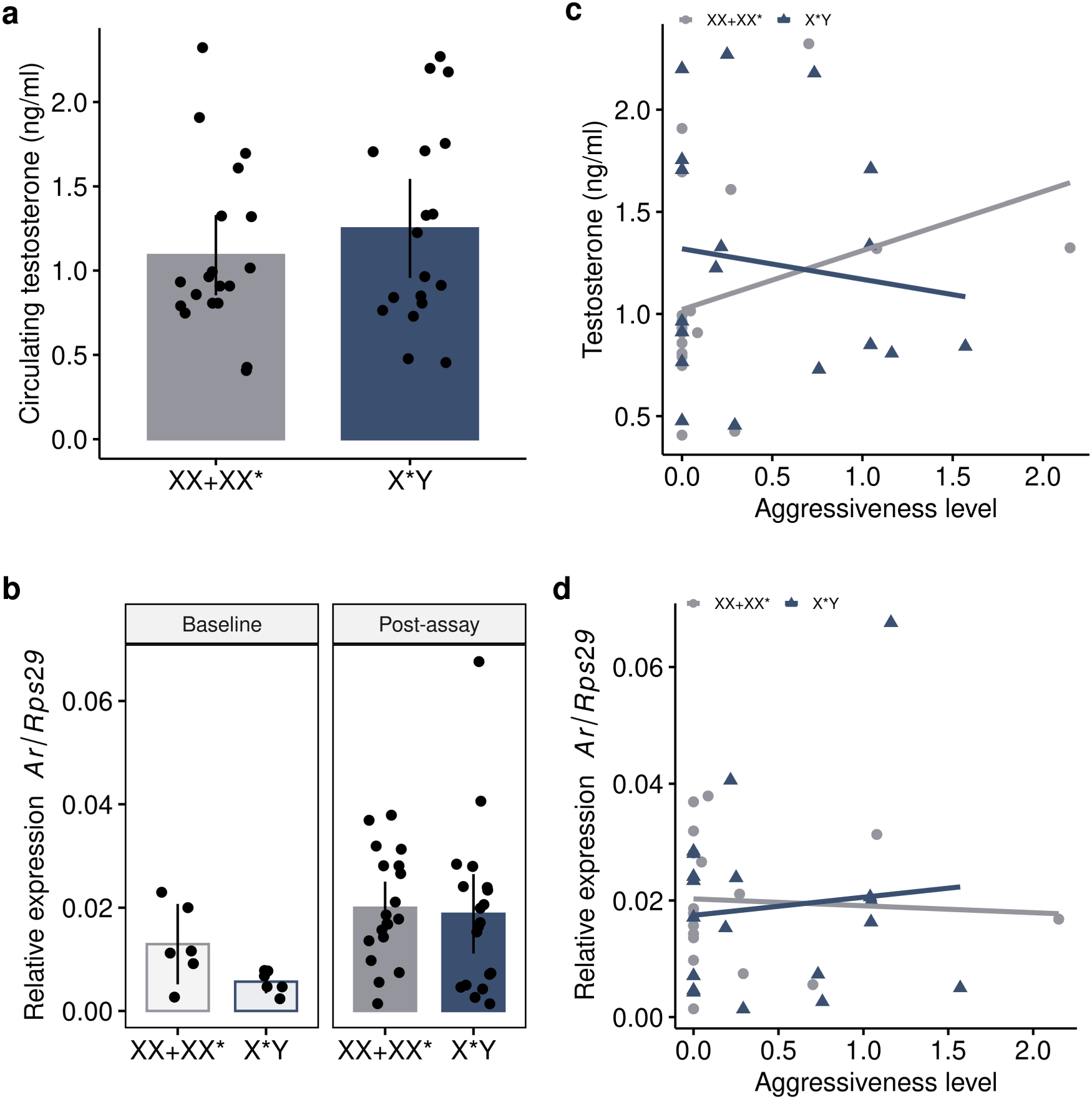
Androgen levels and correlation with aggression in X*Y and XX/XX* females. **a, c)** serum testosterone concentration in X*Y and XX/XX* females following the resident-intruder test and correlation between testosterone levels and the level of aggressiveness, respectively. n_XX+XX*_=19 and n_X*Y_ =18. **b,** relative basal and post-assay expression of the androgen receptor (*Ar*) and **d**) correlation between post-assay *Ar* expression and the level of aggressiveness. Baseline, n_XX+XX*_=6 and n_X*Y_ =6; Post-assay, n_XX+XX*_=19 and n_X*Y_ =19. XX and XX* females formed the control group with low aggressiveness (Saunders et al. 2016). Bars are mean ± 95% confidence intervals and dots/triangles represent raw data. Regression lines are for visual purpose only.

### Sry and the dopaminergic pathway: Th, MaoA and Drd2

The basal expression level of *Sry* in the whole brain of X*Y females is extremely low, with an average at 2.2e-5 [95% CI = 1.38e-5 – 3.06e-5], and with no significant difference between basal and post-assay expression levels (Fig. 3a, LM, df = 23, estimate = –3.2e-6 ± 4.6e-6 SE, p=0.5). Baseline and post-assay *Th* expression levels are not significantly different between X*Y and XX/XX* females, despite a tendency for a greater baseline expression level in XX/XX* females (Fig. 3b, post-hoc Sidak, df = 46, Baseline ratio = 2.86 ± 1.23 SE, p=0.07; Post-assay ratio = 1.13 ± 0.27 SE, p=0.98). However, only X*Y females show higher post-assay *Th* expression levels compared to baseline expression levels (post-hoc Sidak, df = 46, X*Y ratio = 0.32 ± 0.11 SE, p=0.008; XX/XX* ratio= 0.81 ± 0.28 SE, p=0.96). Baseline and post-assay *MaoA* expression levels are lower in X*Y females (Fig. 3c, post-hoc Sidak, df = 46, Baseline ratio =7.05 ± 2.27 SE, p<0.0001; Post-assay ratio = 4.26 ± 0.77 SE, p<0.0001), even though X*Y females show a greater post-assay expression level compared to baseline (post-hoc Sidak, df = 46, ratio = 0.42 ± 0.11 SE, p=0.006). This difference between post-assay and baseline expression levels is not observed in XX/XX* females (post-hoc Sidak, df = 46, ratio= 0.69 ± 0.18 SE, p=0.5). *Drd2* expression levels are not significantly different between females, regardless of whether they were tested (Fig. 3d, post-hoc Sidak, df = 46, Post-assay ratio = 1.17 ± 0.28 SE, p=0.93) or not (post-hoc Sidak, df = 46, Baseline ratio = 2.09 ± 0.89 SE, p =0.31). There is also no significant differences between post-assay and baseline levels for each genotypic group, despite a tendency for a greater post-assay expression level in X*Y females (post-hoc Sidak, df = 46, X*Y ratio= 0.43 ± 0.15 SE, p=0.07; XX/XX* ratio = 0.77 ± 0.27 SE, p= 0.91). Finally, none of these genes have an expression level that significantly correlates with aggression in X*Y (Fig 3e-h, Kendall test, *Sry*: τ=0.16, p =0.84; *Th*: τ= 0.03, p=0.94; *MaoA*: τ=0.01, p =0.94; *Drd2*: τ= 0.07, p=0.94) or XX/XX* females (Kendall test, *Th*: τ= 0.02 p=0.97; *MaoA*: τ=0.05, p =0.697; *Drd2*: τ= 0.007, p=0.97).

**Figure 3.**
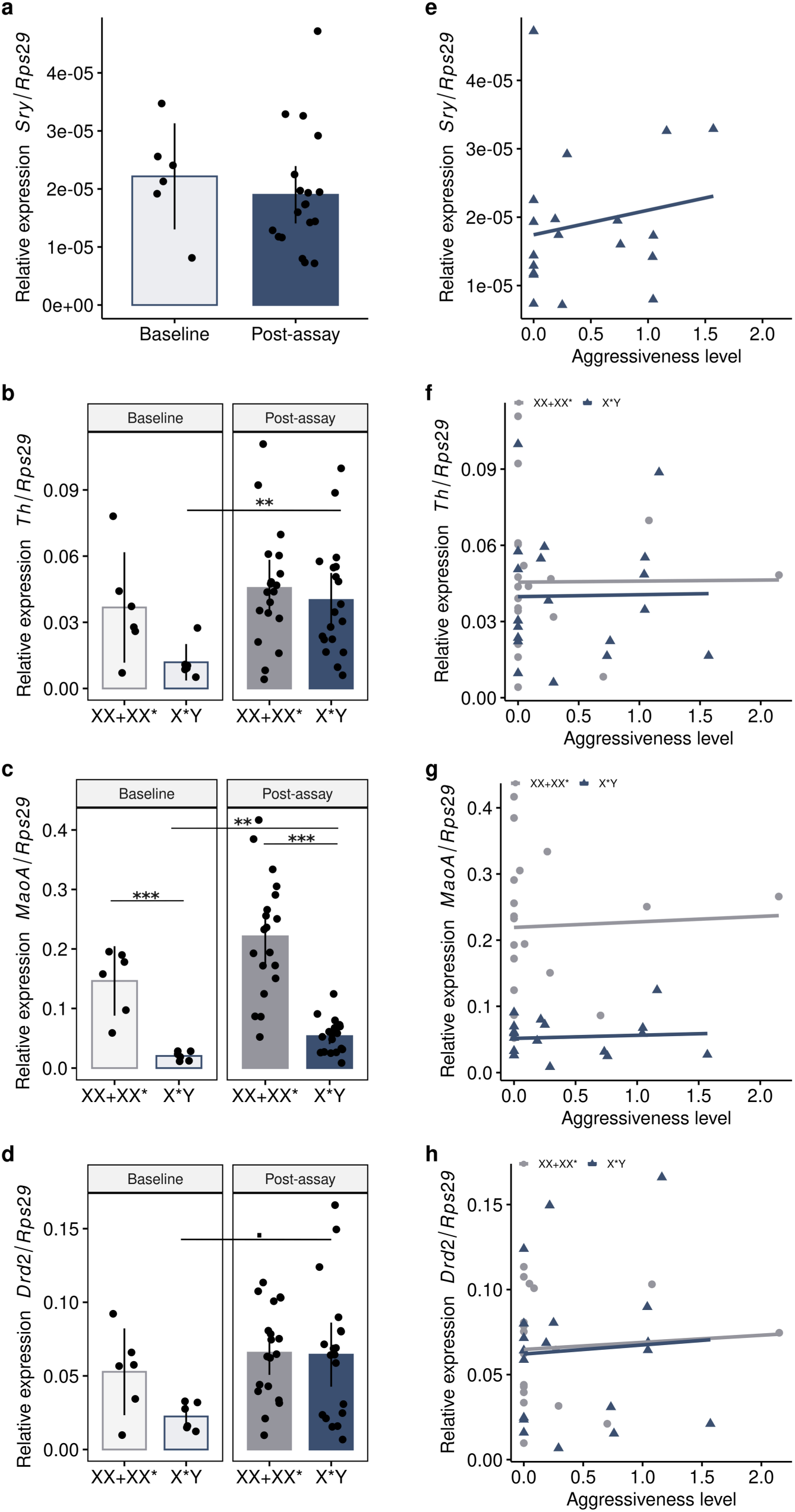
Relative expression of *Sry* and sex-linked genes from the dopaminergic pathway in the whole brain. Relative baseline and post-assay expression of *Sry*. (**a**) *Th* (**b**) *MaoA* (**c**) and *Drd2* (**d**), and correlation between expression levels and aggression (**e**, **f**, **g**, **h** for *Sry, Th*, *MaoA,* and *Drd2,* respectively) in X*Y and XX/XX* females. Bars are mean ± 95% confidence intervals and dots/triangles represent raw data. XX and XX* females formed the control group with low aggressiveness (Saunders et al. 2016). Baseline, n_XX+XX*_=6 and n_X*Y_ =6; Post-assay, n_XX+XX*_=19 and n_X*Y_ =19. Regression lines are for visual purpose only. ***p≤0.001, **p≤0.01, *p≤0.05, ·p ≤0.1

### Neuropeptides: Oxt and Avp

*Avp* expression levels are overall not significantly different between X*Y and XX/XX* females (Fig. 4a, post-hoc Sidak, df = 46, Baseline ratio= 0.88 ± 0.34 SE, p=0.99; Post-assay ratio= 0.86 ± 0.19 SE, p=0.93). Moreover, *Avp* expression are very low, with an average relative post-assay level at 0.0003 [95% CI = 0.0002-0.0004] for instance, whereas the post-assay *Oxt* level is at 0.02 [95% CI= 0.016-0.034] in average for both genotypic group. There is also no significant differences comparing baseline and post-assay expression levels for X*Y (post-hoc Sidak, df = 46, ratio= 0.9 ± 0.28 SE, p=0.99) and XX/XX* (post-hoc Sidak, df = 46, ratio= 0.92 ± 0.29 SE, p=0.99) females, nor there is a significant correlation between aggressiveness and *Avp* levels (Fig. 4b) in each genotypic group (Kendall test, X*Y, τ = 0.22, p = 0.75; XX/XX*, τ=-0.26, p=0.93). Baseline *Oxt* expression levels are significantly greater in XX/XX* than in X*Y females (Fig 4c, Post-hoc Sidak, df = 46, ratio=3.66 ± 1.78 SE, p=0.04) but there is no difference between females looking at post-assay expression levels (Post-hoc Sidak, df = 46, ratio=1.05 ± 0.29 SE, p=0.99). This is consistent with post-assay *Oxt* expression levels which are greater than baseline expression levels in X*Y females only (Post-hoc Sidak, df = 46, X*Y ratio=0.3 ± 0.12 SE, p=0.01; XX/XX* ratio=1.04 ± 0.41 SE, p=1). Finally, we did not find a significant correlation between the level of aggressiveness and *Oxt* expression levels in X*Y (Fig 4d, Kendall test, τ = 0.22, p=0.75), nor in XX/XX* (τ=0.14, p=0.97) females.

**Figure 4.**
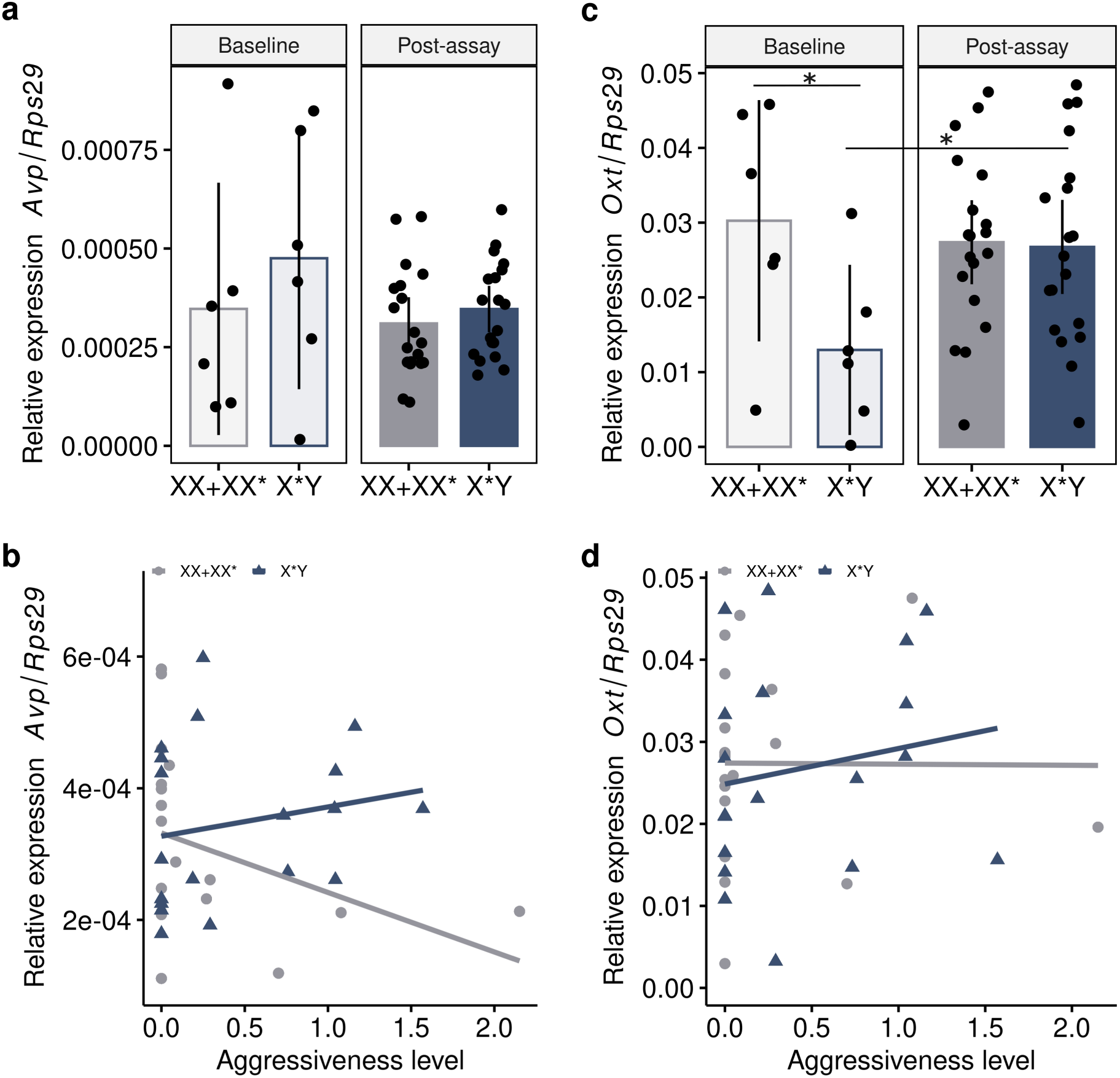
Relative expression of vasopressin (*Avp*) and oxytocin (*Oxt*) in the brain of X*Y and XX/XX* females. Relative baseline and post-assay expression of *Avp*. (**a**) and *Oxt* (**c**) and correlations with aggression type (**b**, *Avp*, and **d**, *Oxt*). Bars are mean ± 95% confidence intervals and dots/triangles represent raw data. XX and XX* females formed the control group with low aggressiveness (Saunders et al. 2016). Baseline, n_XX+XX*_=6 and n_X*Y_ =6; Post-assay, n_XX+XX*_=19 and n_X*Y_ =19. Regression lines are for visual purpose only. ***p≤0.001, **p≤0.01, *p≤0.05, ·p ≤0.1

## Discussion

In the African pygmy mouse, sex-reversed X*Y females have previously been reported to be highly territorial but the molecular basis of this behaviour is unknown. By correlating behaviours with testosterone and candidate gene expression levels, our results suggest that the territoriality of the X*Y females is more likely to be driven by the influence of sex chromosomes on dopamine and/or neuropeptide levels than by the androgen pathway. We acknowledge that these analyses are conducted on whole brain samples, an approach that is likely to dilute signals and limits the detection of region-specific patterns. Nevertheless, given the limited availability of samples within the colony, we believe the differences reported here using the whole tissue provide a valuable foundation and exciting prospects to future in-depth studies targeting specific brain regions.

## Tendencies for enhanced aggressiveness in X*Y females

In a previous study, Saunders et al. (2016) highlighted that X*Y females were more aggressive than both XX and XX* females during a resident-intruder assay with the opposite sex (i.e. males): X*Y females had a shorter latency to first attack and attack approximately three times as much as the other females. Here, our results reveal a reduced level of aggressiveness in X*Y females in comparison to this primary observation. We did not find significant differences in aggressiveness level between X*Y and XX/XX* females, although X*Y females tend to be more offensive and XX/XX* seem less likely to be aggressive (Fig. 1). In addition, our results show that the effect size of aggressive behaviours in X*Y females is not as strong as the one previously reported (in Saunders et al. 2016; see Fig. 1), with a greater variance in aggressiveness level. Because we used individuals from the same colony, it is likely that these differences reflect an artificial selection due to breeding conditions (i.e., ‘breeding facility’ effect). This phenomenon could be common in animals that are held in captivity. In birds for instance, it has been shown that one generation of captive breeding is sufficient for the loss of anti-predatory behaviours (Carrete & Tella, 2015). In our case, there has been a gap of about nine years between experimentations representing 18 generations (given that there are approximately two generations per year in our breeding colony). Considering that highly aggressive X*Y females can be difficult to pair – some were even excluded from the breeding program if they have killed a male (Heitzmann et al. 2023) –, it is likely that highest levels of aggressiveness were artificially selected against in our breeding facility. Consequently, the lack of significance may reflect a combination of lower aggressiveness (i.e., lower effect size), and of a lower statistical power because of a sample size too low to detect smaller effects. These finding highlights that although captivity provide great advantages by enhancing research accessibility, it also has its limitations and can be subject to artificial selection which can bias phenotypic studies. Including animals from the wild would greatly benefits future studies investigating territorial aggression in the African pygmy mouse. Nonetheless, we still observed aggressive females, enabling the establishment of a relationship between aggressiveness and hormonal or gene expression levels.

## No causal link between androgens and territoriality

Gonadal hormones and sex chromosomes interact to shape brain neural circuits underlying behaviours. The role of steroid hormones and especially testosterone have been widely investigated in relation to territorial aggression (e.g. Giammanco et al. 2005; Wu et al. 2009; Xu et al. 2012). Testosterone is a key factor for the masculinization of brain neural pathway, either directly or through the aromatization of testosterone to estrogen (Han & De Vries, 2003; Phoenix, 2009; Phoenix et al. 1959; Wu et al. 2009). More precisely, testosterone has been shown to regulate social aggression in males, but also in females as illustrated in some bird species, rats and humans (Cantarero et al. 2015; Denson et al. 2018; Giammanco et al. 2005; Gill et al. 2007; Rose et al. 1971; Wu et al. 2009). Here, testosterone levels are comparable between females, as previously observed when examining basal testosterone levels (see Veyrunes et al. 2024), albeit concentrations seem overall greater in the present study, which could be explained by the post-behavioural assay context. We did not find a correlation between aggressiveness and testosterone levels, with a slight tendency for a positive relationship in XX/XX* females only, a find that is mainly driven by one outlier individual (Fig. 2c). Because testosterone action depends on its receptor, the androgen receptor (AR) (see Cunningham et al. 2012), we also evaluated the expression of *Ar* and included females that did not perform the test to control for baseline expression levels. It is also noteworthy that the *Ar* gene is on the X chromosome and in the region that does not recombine with the X* chromosome (Saunders et al. unpublished data). Its expression is thus likely to be influenced by differences in the sex chromosome complement between females (e.g., X dosage, divergence between X and X* copies). There are no differences in *Ar* expression levels between females but most importantly, basal and post-assay levels remain within the same expression range, suggesting that resident-intruder encounter does not induce an increase in *Ar* expression. Moreover, there is no apparent correlation between *Ar* expression and aggression. Taken together, our results are not sufficient to support an involvement of androgens in females’ territorial aggression in the African pygmy mouse.

It has been shown that the ratio of corticosterone over testosterone could be a more reliable measure to correlate with aggression than testosterone itself (Montoya et al. 2012; Terburg et al. 2009). Interestingly, X*Y females exhibit lower basal corticosterone than XX and XX* females, but this difference disappears in stressful conditions (Veyrunes et al. 2024), which are presumed to occur here. Given the greater fold-change in corticosterone levels between basal and stressful conditions in X*Y females, it is possible that these females display a different sensitivity to corticosterone. Nevertheless, this suggests that the ratio of testosterone/corticosterone is unlikely to be a better predictor of aggressiveness than testosterone alone in *M. minutoides*. Secondly, it has been shown that the intrauterine hormonal environment could also influence aggressiveness, where females developing between two males are more aggressive than females developing with adjacent females (Ryan & Vandenbergh, 2002). However, in our case, due to a complex meiotic drive (Saunders et al. 2022), the least aggressive XX females have mostly brothers, while the most aggressive X*Y females have mainly sisters, which does not support an intrauterine effect of androgens on the aggressive behaviours of adult females. Finally, estrogens have also been shown to regulate female aggression in mice and humans (Denson et al. 2018; Hashikawa et al. 2017; Woodley & Moore, 1999). While we did not include estrogens here, a recent investigation highlighted no differences in circulating estradiol concentrations between female genotypes of the pygmy mouse (Veyrunes et al. 2024). Nonetheless, future studies looking at estrogen receptors in specific regions of the brain are mandatory to completely rule out the role of estrogens and more generally, of hormones.

## Resident-intruder assays stimulate the dopaminergic pathway in X*Y females only

Dopamine is a catecholamine neurotransmitter that regulates many social behaviours and elevated levels of dopamine have been associated with aggressive behaviours (Czech et al. 2012; Korzan et al. 2006; Mahadevia et al. 2021; Scott et al. 2008). In *M. minutoides*, dopamine levels are likely to vary according to the sex chromosome complement of females (XX, XX* or X*Y). First, the sex-determining *Sry* gene which can promote dopamine synthesis (Dewing et al. 2006) has been shown to be expressed in the brain of X*Y females (Rahmoun et al. 2014). Then, the X-linked *MaoA* gene responsible for dopamine degradation is located in the non-recombining region of X and X* chromosomes, and in multiple copies on the X chromosome only (unpublished data). Finally, dopamine precursor (*Th*), and dopamine receptor D2 (*Drd2) –* which inhibits dopamine release and synthesis in an auto-negative feedback loop *–,* are also sex-linked in *M. minutoides*. *Th* and *Drd2* are located in the recombining regions of two distinct neo-sex chromosomes but because their inheritance pattern differs between sexual genotype, their expression pattern may vary accordingly (Saunders et al. unpublished data; Heitzmann et al. 2025). Among these four genes, *Th* and *MaoA* had previously been identified as potential candidates underlying female genotype divergences in social behaviours (Heitzmann et al. 2023; 2025). A study on the whole-brain transcriptome revealed greater *Th* and lower *MaoA* expression levels in X*Y females compared to the other female genotypes (Heitzmann et al. 2025). Consistently, neuroanatomical investigations of the maternal behaviours circuitry also suggested greater Th levels in X*Y females, although these results were not statistically significant (Heitzmann et al. 2023). In the present study, we confirm the overall lower expression levels of *MaoA* in X*Y females compared to XX and XX* females, but we also observed a lower basal *Th* expression level in X*Y females, contrary to previous findings. Female differences in *Th* expression disappear post-assay whereas *MaoA* expression levels remain significantly lower in X*Y females. Taken together, these results suggest elevated post-assay dopamine levels in X*Y females compared to XX/XX* females. In fact, only X*Y females show greater post-assay expression of *Th*, *MaoA* and presumably *Drd2* when compared to basal expression. These results are in line with a stimulation of dopamine metabolism in X*Y females only, where encounters with unfamiliar males induce dopamine production, which in turn triggers compensatory upregulation of its receptor and degradation pathways to restore homeostasis.

Given the absence of differences in *Sry* expression –which was relatively low compared to previous reports (Rahmoun et al. 2014)– it is unlikely that *Sry* enhances dopamine activity via *Th* upregulation (Dewing et al. 2006). However, since *Sry* expression is restricted to specific brain regions (e.g. Lahr et al. 1995), its potential involvement cannot be entirely ruled out, as its signal may be diluted in whole-brain analyses. Taken together, our results suggest that the dopaminergic pathway is stimulated following an agonistic assay in X*Y females only. Nevertheless, because there is no apparent correlation between aggressiveness and gene expression levels, this highlights that aggression is a complex and multifactorial behaviour that cannot be reduced to a few genes. While dopamine may be permissive for aggression, it does not necessarily lead to it; the behavioural output and onset of aggression seem to vary according to other factors. In the future, multifactorial investigations as well as quantification of dopamine levels in brain regions of interest (e.g., ventral tegmental area, nucleus accumbens; Mahadevia et al. 2021) should help unravel the proximate causes of territorial aggression in the African pygmy mouse.

## Vasopressin and oxytocin

AVP and OXT are two neuropeptides involved in the regulation of social behaviours including aggression (e.g. Gatewood et al. 2006; Moura Oliveira et al. 2021). While both are known to stimulate maternal aggression (see Bosch & Neumann, 2012), their influence on territorial aggression is rather species-dependant due to controversial results with both stimulating and inhibitory effects (e.g. Bester-Meredith & Marler; de Moura Oliveira et al. 2021; Gatewood et al. 2006; Tan et al. 2019; Everts et al. 1997). For instance, AVP immunoreactive density of fibers correlates positively with aggression in the California mouse, *Peromyscus californicus*, while *Avp* inhibits aggression in rats (Bester-Meredith & Marler; de Moura Oliveira et al. 2021; Everts et al. 1997). Here, our results do not support an increase in *Avp* expression following the resident-intruder assay, which is moreover very low compared to *Oxt* levels for instance (Fig 4a). We also did not find a correlation between *Avp* and aggression levels in females (Fig4b), despite a steeper negative slope in XX/XX* females. Given the low levels and narrow range of expression, we assume it to reflect an artefact rather than a true but undetected effect. It is thus unlikely that *Avp* influences territorial aggression in X*Y females or XX/XX* females. Conversely, our data support an increase in *Oxt* expression following the resident-intruder assay in X*Y females only, as observed for dopamine. Given the well-documented interactions between oxytocin and dopamine systems in shaping social behaviours (Baskerville & Douglas, 2010; Melis et al., 2007; Scott et al., 2015; Peris et al. 2017), these findings suggest that their interplay may underlie the unique behavioural phenotype of X*Y females. An interesting possibility is that *Oxt* increase reflects how females cope with the stress induced by the resident-intruder experiment. In the violet-eared waxbill for instance, Oxt has been hypothesized to enable (and not trigger) aggression due to its anxiolytic effect, as aggressive individuals tend to be less responsive to stress (Goodson et al. 2015). Therefore, one might suggest that O*xt* and dopamine interact in stressful conditions to shape (at least partly) X*Y female behaviours during female-male encounters, which remains to be tested.

## Conclusion

X*Y females of the African pygmy mouse have previously been shown to be highly territorial compared to other females. We highlight that the dopaminergic and Avp/oxytocin pathways, rather than androgens, are more likely to drive this trait in *M. minutoides*. Because the increase in dopamine/*Oxt* levels does not correlate with agonistic behaviours, the motivation to be aggressive may thus depend on other interacting factors. In addition, because changes in gene expression mostly occur in X*Y females, it is likely that X*Y and XX/XX* females have different proximate mechanisms promoting aggression. Future investigations such as *in situ* hybridization in specific regions of the brain (e.g. ventral tegmental area) that include wild animals to remove breeding effects and other biological factors should help unravel the proximate causes of territorial aggression. Nonetheless, our findings strongly suggest differences in activation of the dopaminergic pathway between X*Y and XX/XX* females. Given that this pathway is tightly linked to sex and neo-sex chromosomes in *M. minutoides*, our findings support an influence of sex reversal on the dopaminergic system and therefore a direct influence of sex chromosomes on the sexual differentiation of the brain. This further challenges the paradigm that has dominated for decades, of the predominant, if not exclusive role of hormones (see Breedlove et al. 1999; Arnold, 2009).

## Data availability

Script and data will be available on Zenodo (doi; 10.5281/zenodo.17834305) upon publication.

## Acknowledgements

We thank the CECEMA for the housing of animals and Marie Challe for her assistance on the maintenance of the breeding colony. We also thank the MGX-qPCR Platform for technical support and Philippe Clair for his assistance on RT-qPCR as well as on data analysis on LightCycler 480 software. We thank Emmanuel Valjent and Francis Poulat for constructive comments and discussions on the topic. We kindly thank Phillip C. Boan for proofreading the paper. A preprint version of this article has been peer-reviewed and recommended by PCI Evol Biol (“https://doi.org/10.24072/pci.evolbiol.100946”). We thank two anonymous reviewer and Barbara Class (data editor) for their careful evaluation and comments. This study was supported by the ANR grant SEXREV (no. 18-CE02-0018-01).

